# BRAIN-DERIVED NEUROTROPHIC FACTOR IN AN ORBITOFRONTAL CORTICAL-DORSOLATERAL STRIATAL CIRCUIT GATES ALCOHOL CONSUMPTION

**DOI:** 10.1101/2021.09.10.459813

**Authors:** Jeffrey J. Moffat, Samuel A. Sakhai, Yann Ehinger, Khanhky Phamluong, Dorit Ron

## Abstract

Brain-derived neurotrophic factor (BDNF) signaling in the dorsolateral striatum (DLS) gates alcohol self-administration in rodents. The major source of BDNF in the striatum is the cortex, and we recently found that BDNF-expressing neurons in the ventrolateral orbitofrontal cortex (vlOFC) extend axonal projections to the DLS. We therefore hypothesized that BDNF in the vlOFC to DLS circuit moderates alcohol intake. We show that overexpression of BDNF in the vlOFC, which activates BDNF signaling in the DLS, is sufficient to attenuate voluntary consumption and seeking of 20% alcohol in the home cage using a two-bottle choice paradigm. Overexpressing BDNF in the vlOFC had no effect on the consumption of a sweetened saccharin solution. In addition, BDNF overexpression in the neighboring motor cortex did not alter alcohol intake. Finally, pathway-specific overexpression of BDNF in DLS-projecting vlOFC neurons significantly reduced alcohol intake and preference. Overall, BDNF in the vlOFC, and specifically in a vlOFC-DLS pathway, keeps alcohol drinking in moderation.

## Introduction

Alcohol use disorder (AUD) is characterized by compulsive alcohol use despite negative consequences [1,2]. Although more than 50% of adults in the United States regularly consume alcohol, only 6.2% can be considered heavy drinkers [2,3], and the vast majority of alcohol drinkers do not lose control over their alcohol intake [1,2]. Studies from our lab and others suggest that endogenous factors protect the majority of alcohol users from developing AUDs by keeping alcohol consumption in moderation [4-6].

Among these endogenous protective factors is brain-derived neurotrophic factor (BDNF) [4,5]. BDNF is a member of the nerve growth factor (NGF) family of neurotrophic factors and is highly expressed in the central nervous system (CNS) [7-9]. Over the past decade, we and others observed that BDNF interacts with alcohol in a unique way [4,5]. Specifically, we found that activation of BDNF signaling in the dorsolateral striatum (DLS), which is mediated via the receptor tyrosine kinase, tropomyosin-related kinase B (TrkB), keeps alcohol intake in moderation [10-13]. We further showed that escalation of drinking occurs when this protective signaling pathway ceases to function [14-17].

The majority of BDNF is stored in presynaptic dense core vesicles and is released upon neuronal depolarization [18-21]. This suggests that BDNF which activates TrkB signaling in the DLS originates from neurons in a different brain region. Indeed, previous studies show that the primary source of BDNF in the adult striatum is the cerebral cortex [22-24]. We recently surveyed BDNF-positive cortical inputs into the DLS and, surprisingly, found that BDNF-expressing neurons in the ventrolateral orbitofrontal cortex (vlOFC) extend axonal projections to the DLS [25]. We also showed that overexpression of BDNF in the vlOFC activates TrkB signaling in the DLS and not in the adjacent dorsomedial striatum (DMS) [25]. We therefore hypothesized that BDNF released from vlOFC neurons into the DLS acts to keep alcohol intake in moderation.

## Methods and Materials

### Reagents

Mouse anti-NeuN antibody (MAB377) was obtained from Millipore (Billerica, MA). Anti-GFP antibody (Cat #AB290) was purchased from Abcam (Cambridge, UK). Donkey anti-mouse IgG AlexaFluor 564, donkey anti-chicken AlexaFluor 488 antibodies, were purchased from Life Technologies (Grand Island, NY). Ethyl alcohol (190 proof) was purchased from VWR (Radnor, PA). Saccharin (saccharin sodium salt hydrate) was purchased from Sigma Aldrich (St. Louis, MO). Other common reagents were from Sigma Aldrich (St. Louis, MO) or Fisher Scientific (Pittsburgh, PA).

### Animals

Male C57/BL6 and CamKIIα-Cre mice (6-8 weeks old at time of purchase) were obtained from Jackson Labs (Bar Harbor, Maine). Mice were individually housed on paper-chip bedding (Teklad #7084), under a reverse 12-hour light-dark cycle (lights on 10 am to 10 pm). Temperature was kept constant at 22 ± 2°C and relative humidity was maintained at 50 ± 5%. Mice were allowed access to food (Teklad Global Diet #2918) and tap water *ad libitum*. All animal procedures were approved by the University of California, San Francisco Institutional Animal Care and Use Committee and were conducted in agreement with the Association for Assessment and Accreditation of Laboratory Animal Care (AAALAC, UCSF).

### Virus production

AAV1/2-IRES-CMV-BDNF-GFP (AAV-BDNF; 1 × 10^12^ gc/ml) and AAV1/2-IRES-CMV-GFP (AAV-GFP; 1 × 10^12^ gc/ml) were designed as described in [16] and were purchased from the University of North Carolina Viral Vector Core. AAV2-CMV-GFP-mBDNF (AAV2-BDNF; 7.7 × 10^12^gc/ml) and AAV2-CMV-GFP (AAV2-GFP; 1 × 10^13^ gc/ml) were purchased from Vector Biolabs (Malverna, PA). AAV-DIO-Ef1a-BDNF-IRES-mCherry (AAV-DIO-BDNF-mCherry; 1 × 10^12^ gc/ml) and AAV-DIO-Ef1a – mCherry (AAV-DIO-mCherry; 1 × 10^12^ gc/ml) were constructed in conjunction with C&M Biolabs (Richmond, CA) and were purchased from the University of Pennsylvania Viral Vector Core. AAV5.CMV.HI.eGFP-Cre.WPRE.SV40 (AAV-Cre; 4.8 × 10^12^ gc/ml) was also purchased from the University of Pennsylvania Viral Vector Core.

### Stereotaxic viral infection

Animals underwent stereotaxic surgery targeting the OFC, or motor cortex and/or the DLS. Mice were anesthetized by vaporized isoflurane and were placed in a digital stereotaxic frame (David Kopf Instruments, Tujunga, CA). Two holes were drilled above the site of viral injection. The injectors (stainless tubing, 33 gauges; Small Parts Incorporated, Logansport, IN) were slowly lowered into the target region. The injectors were connected to Hamilton syringes (10 µl; 1701, Harvard Apparatus, Holliston, MA), and the infusion was controlled by an automatic pump at a rate of 0.1 µl/min (Harvard Apparatus, Holliston, MA). The injectors remained in place for an additional 10 minutes to allow the virus to diffuse, and then were slowly removed. Mice were allowed to recover in their home cage for at least one week before further testing.

To overexpress BDNF in the vlOFC, animals were infused with 1 µl of AAV-BDNF (7.7 × 10^12^gc/ml) or AAV-GFP (1 × 10^13^ gc/ml) per injection site, with two injections per hemisphere (AP: +3 and +2.58, ML: ± 1.2, DV: -2.3, infusion at -2.2 and -2.55 and -2.5 from bregma for each injection respectively).

To overexpress BDNF in the motor cortex, animals were infused with 1 µl of AAV-BDNF (1 × 10^12^ gc/ml) or AAV-GFP (1 × 10^12^ gc/ml) per hemisphere (AP: +2.58, ML: ± 1.3, DV: -1.35, infusion at -1.15 from bregma).

To confirm the efficacy of our AAV-DIO-BDNF construct, CamKIIα-Cre mice received 1 µl of AAV-DIO BDNF (1 × 10^12^ gc/ml) or AAV-DIO-mCherry (1 × 10^12^ gc/ml) in the vlOFC (AP: +2.58, ML: ± 1.2, DV: -2.55, infusion at -2.5 from bregma). For confirmation of circuit-specific viral targeting, C57Bl6/J mice received 1 µl of AAV-DIO BDNF (1 × 10^12^ gc/ml) in the vlOFC (AP: +2.58, ML: ± 1.2, DV: -2.55, infusion at -2.5 from bregma) and 0.5 µl of AAV-Cre (4.8 × 10^12^ gc/ml) in the DLS (AP: +1.1, ML: ± 2, DV: -2.85, infusion at -2.8 from bregma).

For circuit specific expression of BDNF in the OFC-DLS pathway, mice received 1 µl of AAV-DIO-BDNF (1 × 10^12^ gc/ml) per hemisphere targeting the vlOFC (AP: +2.58, ML: ± 1.2, DV: -2.55, infusion at -2.5 from bregma) and 0.5 µl of AAV-Cre (4.8 × 10^12^ gc/ml) targeting the DLS (AP: +1.1, ML: ± 2, DV: -2.85, infusion at -2.8 from bregma). Control animals received 1 µl of empty vector AAV-DIO-mCherry (1 × 10^12^ gc/ml) in the OFC and 0.5 µl of AAV-Cre (4.8 × 10^12^ gc/ml) in the DLS.

### Solution Preparations

Alcohol solution was prepared from absolute anhydrous alcohol (190 proof) diluted to 20% alcohol (v/v) in tap water. Saccharin solution was diluted to 0.01% or 0.03% saccharin (v/v) in tap water.

### Intermittent Access to 20% Alcohol Two-Bottle Choice

Intermittent access to 20% alcohol, two-bottle choice paradigm (IA20%-2BC) was conducted as previously described [16]. Briefly, mice were given one bottle of 20% alcohol (v/v) and one bottle of water two hours following lights off on Monday, Wednesday, and Friday, with 24 or 48 hours (weekend) of alcohol-deprivation, in which mice consumed only water, in between. The placement of water or alcohol solutions was alternated between each session to control for side preference. Alcohol and water bottles were weighed at the beginning and end of each alcohol drinking session and water and alcohol intake (g/kg) were calculated using weekly animal weights. Corrections were made to account for spillage based on bottles affixed to an empty cage. Preference was calculated by dividing the volume of alcohol consumed by the total liquid intake volume (water + alcohol).

### Home Cage Alcohol Seeking

After the conclusion of IA20%-2BC, one empty alcohol bottle and one water bottle were placed on cages two hours into the dark cycle (following the same pattern used during IA20%-2BC). Contacts with the bottle sipper tubes were detected with lickometers and recorded using Ethovision XT software (Noldus, Leesburg, VA) during the first two minutes of bottle availability. Time licking the empty alcohol bottle and the water bottle were recorded. “Lick time ratios” were graphed as the relative amount of time mice spent licking a particular bottle in one-second time bins. Latency to first and last licks were also recorded.

### Intermittent Access to 0.01 and 0.03% Saccharin Two-Bottle Choice

Saccharin intake was conducted similar to what has been described previously [25]. Animals received one bottle of 0.01% saccharin and one bottle of water continuously for a one-week period. After one week, animals were given 0.03% saccharin solution for an additional week. Saccharin solution drinking (ml/kg of body weight), total fluid intake (ml/kg of body weight) and the preference ratio (volume of saccharin solution intake/total volume of fluid intake) were recorded every 24 hours for the duration of saccharin access. Corrections were made to account for spillage based on bottles affixed to an empty cage.

### Confirmation of viral expression

To confirm viral expression and placement, animals were euthanized by cervical dislocation, and the brains were removed. Brain regions were isolated from a 1-mm-thick coronal section and dissected on ice. Viral infusion into the OFC or DLS was visualized using an EVOS FL tabletop fluorescent microscope (ThermoFisher; Waltham, MA).

### Immunohistochemistry and Microscopy

Following intraperitoneal (IP) administration of euthasol (200 mg/kg), mice were transcardially perfused with phosphate buffered saline, followed by 4% paraformaldehyde (PFA), pH 7.4. Brains were quickly removed post-perfusion and allowed to post-fix for 24 hours prior to cryoprotection in 30% sucrose solution for 3 days at 4 °C. Brains were then sectioned to 50 µm by cryostat (CM3050, Leica, Buffalo Grove, IL), and collected serially 1:6 in 6-well plates and stored in PBS with 0.01% sodium azide. Free-floating PFA-fixed sections were selected, permeablized, and blocked in PBS containing 0.3% Triton and 5% donkey serum for 4 hours. Sections were then incubated for 18 hours at 4°C on an orbital shaker with anti-NeuN antibodies (1:500) and/or anti-GFP (1:5000) diluted in PBS, 3% BSA. Next, sections were washed in PBS then incubated for 4 h with Alexa Fluor 488-labeled donkey anti-rabbit secondary antibody and/or AlexaFluor 564 donkey anti-chicken secondary antibody, diluted in PBS/3% BSA. After staining, sections were rinsed in PBS and cover slipped using Prolong Gold mounting medium. Images were acquired using a Zeiss LSM510 Meta confocal microscope (Zeiss MicroImaging, Jena, Germany) under manufacturer-recommended filter configurations.

### Data Analysis

Data were analyzed with two-way ANOVA, with or without repeated measures and student’s t-test where appropriate. ANOVA and RM ANOVA were used for the two-bottle choice experiments in which the drinking session was the within-subject variable and treatment (BDNF viral overexpression or control) as the between-subject variable. Significant main effects or interactions were further investigated using Bonferroni’s multiple comparisons test. All analyses were run as two-sided and statistical tests were run on Graphpad Prism 8 with a p value of 0.05 considered significant.

## Results

### BDNF in the vlOFC keeps alcohol consumption in moderation

We first sought to determine the influence of BDNF originating from the vlOFC on regulating alcohol drinking. Both hemispheres of the vlOFC of mice were infected with an adeno-associated virus (AAV) expressing BDNF (AAV-BDNF) or control AAV-GFP (**Figure 1a-b**). After one week of recovery, mice were subjected to an intermittent access to 20% alcohol, two-bottle choice (IA20%-2BC) paradigm, in which mice had access to one bottle of 20% alcohol and one bottle of water every other day (**Timeline, Figure 1a)**. We found that mean alcohol consumption and preference were significantly lower in mice overexpressing BDNF in the vlOFC versus GFP expressing mice (**Figure 1c-d**). Water consumption and total fluid consumption were not significantly altered by viral infusion (**Supplemental Figure 1**). Together, our data indicate that BDNF in the vlOFC gates the level of alcohol intake and preference and prevents escalation of alcohol use.

**Figure 1.**
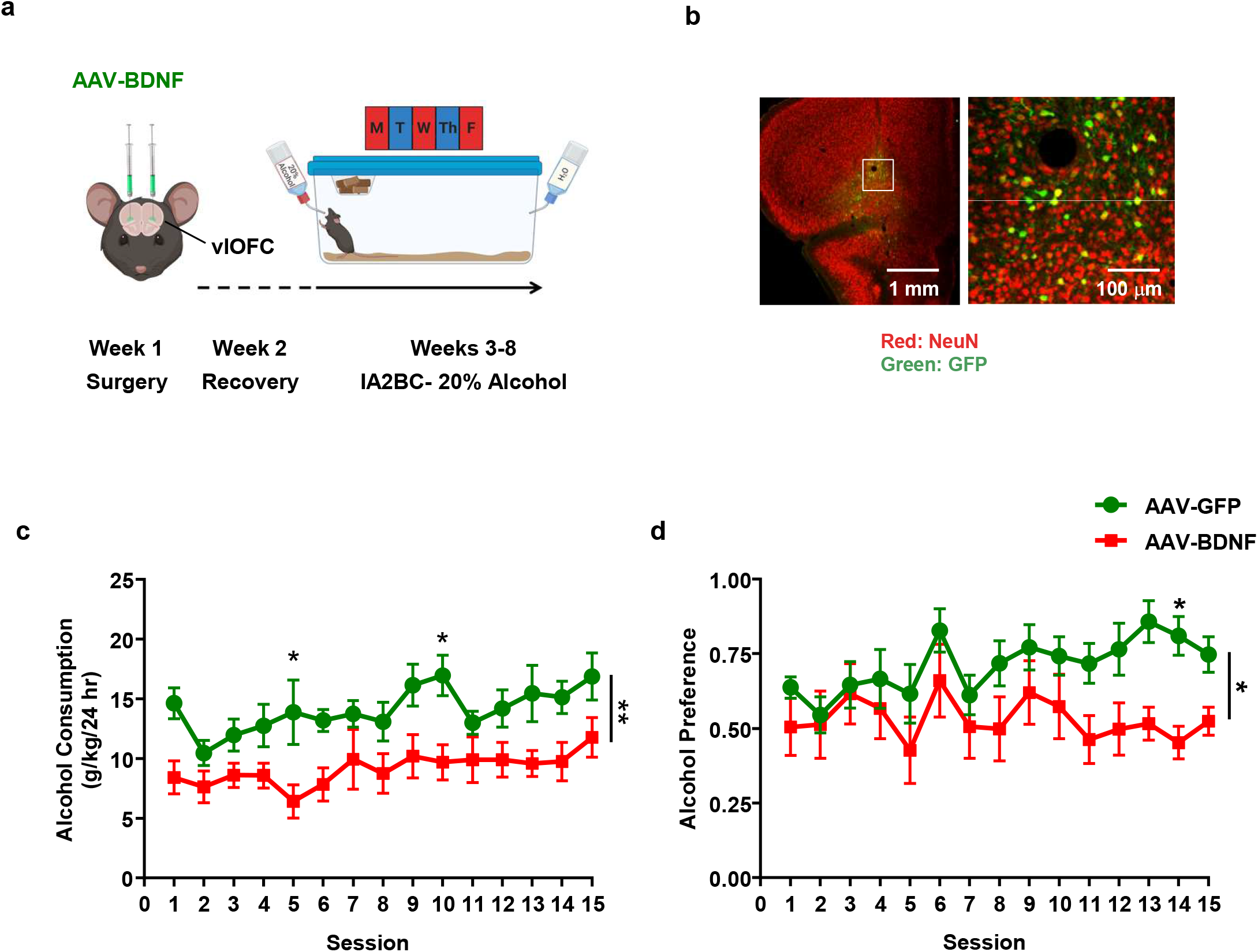
BDNF overexpression in the vlOFC keeps alcohol drinking in moderation. (**a**) Timeline of experiments: Mice received stereotaxic injections of either AAV-BDNF-GFP or AAV-GFP during the first week. Following one week of recovery, mice underwent 15 sessions of IA20%-2BC in the home cage. Alcohol intake was measure at the 24 hour time point (**b**) Confirmation of viral targeting of AAV-BDNF in the vlOFC. Left image is a 5X magnification image of the prefrontal cortex. Right panel is a 20X magnification image of the area of the image in the left panel outlined in white. (**c**) Animals which received AAV-BDNF in the vlOFC maintained moderate levels of alcohol intake as compared to mice infected with AAV-GFP. Two-Way RM ANOVA, main effect of BDNF overexpression, F (1, 18) = 9.712, ^**^p < 0.01, main effect of session, F (14, 252) = 2.835, ^***^p < 0.001, no main effect of interaction. Bonferroni post-hoc testing indicates a significant difference between conditions on sessions 5 and 10, ^*^p < 0.0. (**d**) AAV-BDNF mice exhibited a lower preference for alcohol over water as compared to mice infected with AAV-GFP. Alcohol preference was calculated as the ratio of alcohol intake to total fluid intake at the end of the 24-hour time point. Two-Way RM ANOVA, main effect of BDNF overexpression, F (1, 18) = 7.007, ^*^p < 0.05, no main effect of session or interaction. Bonferroni post-hoc testing indicates a significant difference between conditions on session 14, *p < 0.05. AAV-BDNF n = 10, AAV-GFP n = 10.

### Overexpressing BDNF in the vlOFC reduces alcohol seeking

Following BDNF overexpression and home cage alcohol drinking, as outlined in Figure 1, alcohol seeking behavior was assessed by measuring the amount of time mice spent licking an empty alcohol bottle or a full water bottle immediately after they were affixed to their cage. In the first two minutes of bottle availability, mice that received the AAV-BDNF virus in the vlOFC spent significantly less time licking the empty alcohol bottle than mice that received the AAV-GFP virus **(Figure 2a-b)**. There was no significant difference between the groups in the amount of time spent licking the water bottle **(Supplemental Figure 2a-b)**. There was also no significant difference in the latency to the first or last lick of the alcohol bottle between AAV-BDNF and AAV-GFP groups **(Figure 2c-d)**. Interestingly, mice overexpressing BDNF in the vlOFC displayed a significantly longer latency to their first lick of the water bottle, with 40% of animals never making contact with the bottle at all **(Supplemental Figure 2c)**. BDNF-overexpressing mice also displayed a significantly shorter latency to last lick of the water bottle **(Supplemental Figure 2d)**. This may be due to a lack of interest in newly-placed bottles overall in mice that received AAV-BDNF in the vlOFC. Altogether, these data demonstrate that BDNF overexpression in the vlOFC suppresses alcohol seeking in the home cage environment.

**Figure 2.**
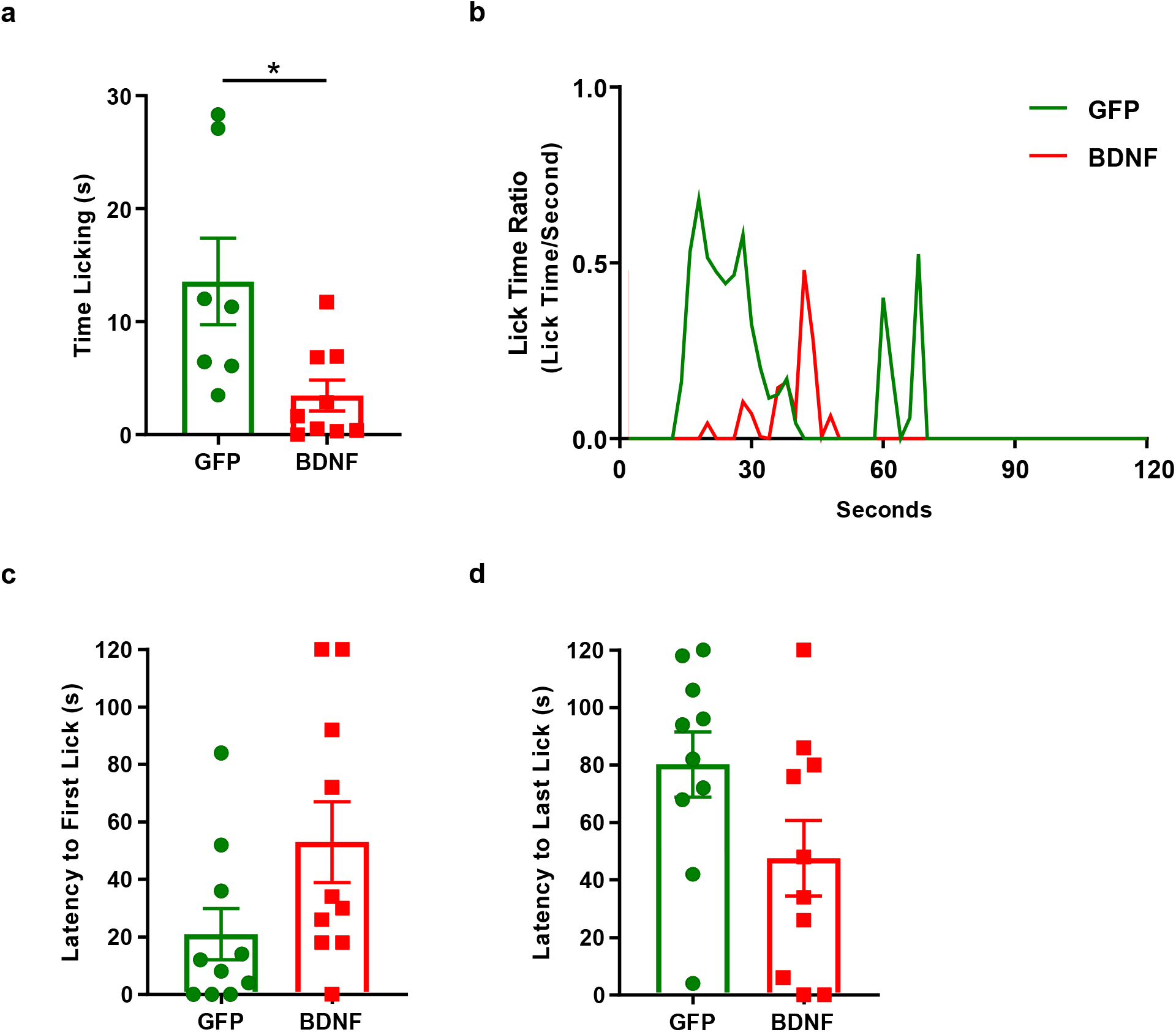
BDNF overexpression in the vlOFC reduces alcohol seeking. (**a-b**) Mice that received AAV-BDNF in the vlOFC and underwent 5 weeks of IA20%-2BC spent significantly less time licking an empty alcohol bottle in the home cages during the first 2 minutes of exposure, compared with mice that received AAV-GFP in the vlOFC. (**a**) AAV-BDNF mice spent significantly less time licking an empty alcohol bottle during a two minute period. Mean time licking is presented with individual data points representing individual animals. Unpaired t Test, t = 2.728, df = 14, *p (two-tailed) < 0.05. (**b**) Representative traces of a single AAV-BDNF infected mouse and a single AAV-GFP infected mouse’s empty alcohol bottle lick time ratio (the amount of time each mouse spent in contact with the bottle sipper tube per second) during a two-minute alcohol seeking test. (**c**) Latency to the first lick of the empty alcohol bottle. Data is presented as the mean amount of time it took for mice from each group to initiate licking of the empty alcohol bottle (maximum value = 120 seconds). Unpaired t Test, t = 1.928, df = 18, p (two-tailed) = 0.0698. (**d**) Latency to last lick of the empty alcohol bottle. Data is presented as the mean amount of time from the final lick of the empty alcohol bottle to the end of the session for mice from each group (maximum value = 120 seconds). Unpaired t Test, t = 1.879, df = 18, p (two-tailed) = 0.0765. AAV-BDNF n = 10, AAV-GFP n = 10.

### BDNF overexpression does not alter saccharin intake

Next, we evaluated whether BDNF’s action on consummatory behavior was specific for alcohol or was shared with other rewarding substances, such as a sweetened saccharin solution. The vlOFC of mice was infused with AAV-BDNF or AAV-GFP one week prior to mice being given continual access to first 0.01% saccharin and water for 1 week, and then 0.03% saccharin solutions and water for another week (**Timeline, Figure 3A**). There were no differences between treatment groups in 0.01% or 0.03% saccharin consumption per day in BDNF-overexpressing mice, compared with GFP controls (**Figure 3B**). Mice overexpressing BDNF in the vlOFC also exhibited no changes in saccharin preference (**Figure 3C**), demonstrating that BDNF in the vlOFC does not impact consumption of a non-alcohol reward.

**Figure 3.**
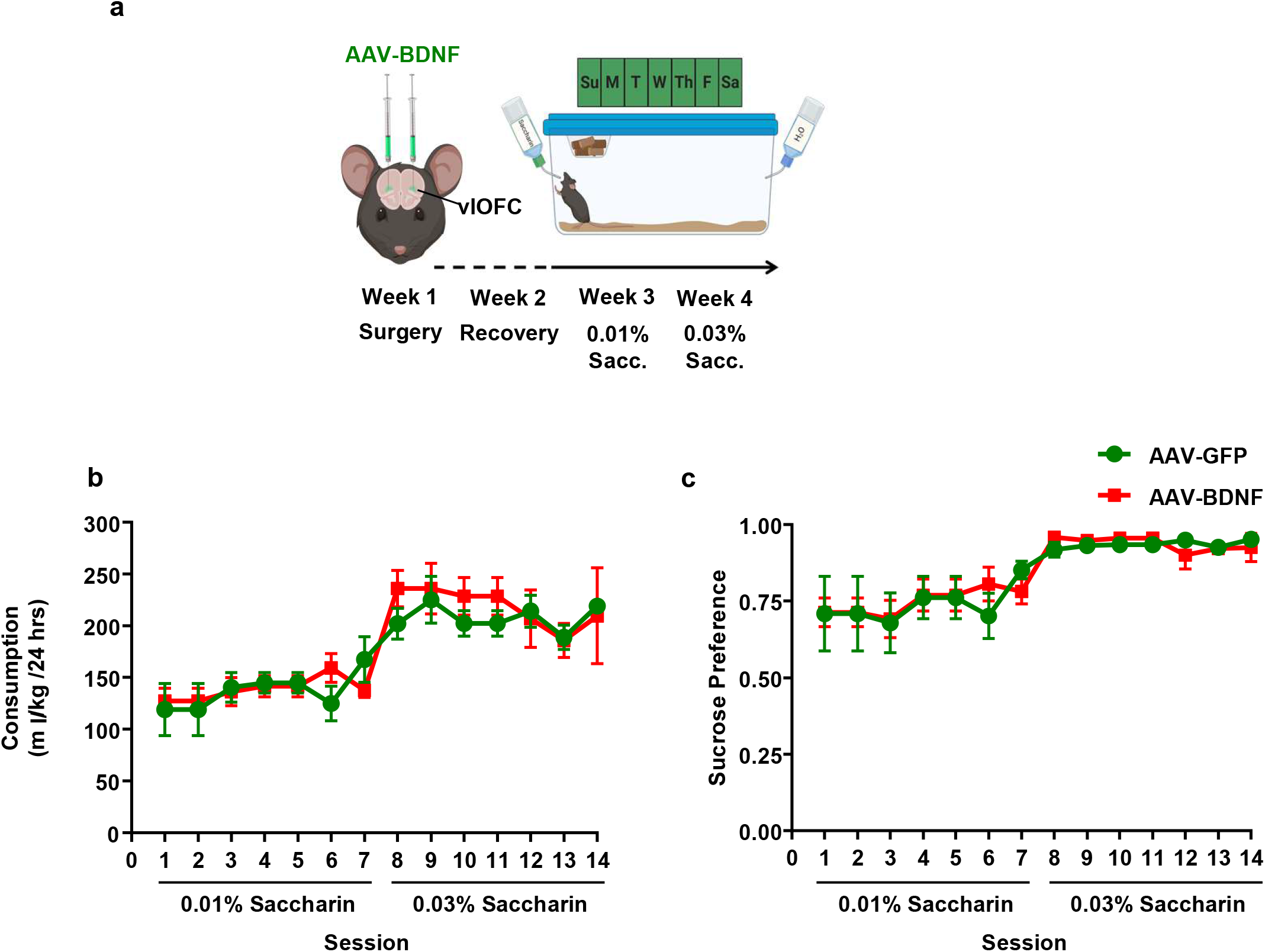
BDNF overexpression in the vlOFC does not alter saccharin consumption. (**a**) Timeline of experiments: AAV-BDNF or AAV-GFP was infused into the vlOFC during week one. After one week of recovery, animals were given a choice of 0.01% saccharin or water followed by the consumption of 0.03% saccharin or water. Animals were given continuous access to each concentration of saccharin solution for a one week period, and saccharin consumption and preference was measured every 24 hours. (**b**) There were no observed differences between groups in saccharin intake (Two-Way ANOVA, no main effect of BDNF overexpression, p > 0.05, or interaction, p > 0.05. Main effect of saccharin concentration, F_(2,24)_ =20.56, ^****^p < 0.0001) or (**c**) preference (Two-Way ANOVA, no main effect of BDNF overexpression, p > 0.05, or interaction, p > 0.05. Main effect of saccharin concentration, F_(1,16)_ =10.64, ^**^p < 0.01). AAV-BDNF n = 5, AAV-GFP n = 5.

### Overexpression of BDNF in the motor cortex does not alter alcohol consumption

Because we previously detected a high number of BDNF-positive neurons in the motor cortex and showed that the DLS also receives BDNF-positive motor-cortical axonal inputs [25], we tested whether BDNF overexpression in the motor cortex would likewise affect alcohol consumption. One week before beginning IA20%-2BC, neurons in the motor cortex were infected with AAV-BDNF or AAV-GFP (**Figure 4a-b**), and alcohol and water consumption were measured after each session (**Timeline Figure 4a**). Alcohol intake and preference were similar in mice overexpressing BDNF and GFP controls over the 12 sessions (**Figure 4c**). In addition, we detected no differences in water or total fluid consumption between groups (**Supplemental Figure 3**). These results indicate that BDNF neurons originating from the motor cortex do not play a role in gating alcohol intake.

**Figure 4.**
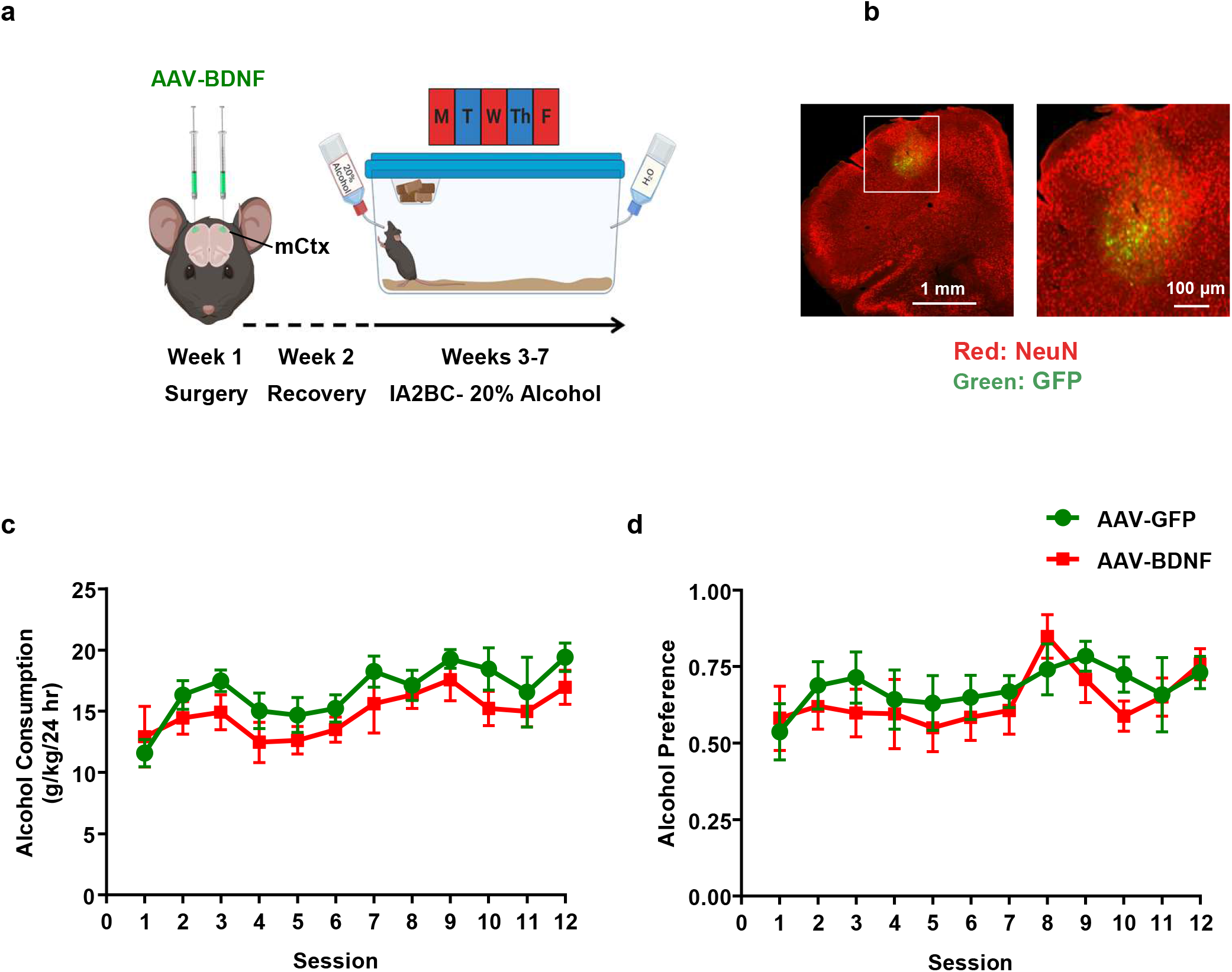
Overexpression of BDNF in the motor cortex does not alter alcohol intake. (**a**) Timeline of experiments: animals were stereotaxically infused with AAV-BDNF virus or AAV-GFP in the motor cortex (mCtx). One week after surgery, mice underwent 12 sessions of IA20%-2BC paradigm and alcohol intake was recorded at the 24 hour time point. (**b**) Confirmation of viral targeting and GFP fluorescence in the mCtx. Left image is a 5X magnification image of the prefrontal cortex. Right panel is a 20X magnification image of the area of the image in the left panel outlined in white. (**c**) Animals which received AAV-BDNF or AAV-GFP did not significantly differ in levels of alcohol consumption (Two-Way RM ANOVA, no main effect of BDNF overexpression, p > 0.05, or interaction, p > 0.05, main effect of session F_(11,132)_ =3.83, ^****^p < 0.0001) or preference (**d**) (Two-Way RM ANOVA, no main effect of BDNF overexpression, p > 0.05, or interaction, p > 0.05, main effect of session F_(11,132)_ =2.21, ^*^p = 0.02). AAV-BDNF n = 7, AAV-GFP n = 7.

### BDNF in vlOFC to DLS circuit regulates alcohol intake

BDNF neurons in the vlOFC project specifically to the DLS [25]. Since overexpressing BDNF in the vlOFC activates TrkB signaling in the DLS [25], and since BDNF-TrkB signaling in the DLS gates alcohol consumption [10-13], we hypothesized that BDNF in vlOFC neurons that specifically project to the DLS gates alcohol drinking. To test this possibility, we utilized an AAV-DIO-BDNF-mCherry construct, which leads to overexpression of BDNF in cells co-expressing Cre recombinase (**Figure 5a**).

**Figure 5.**
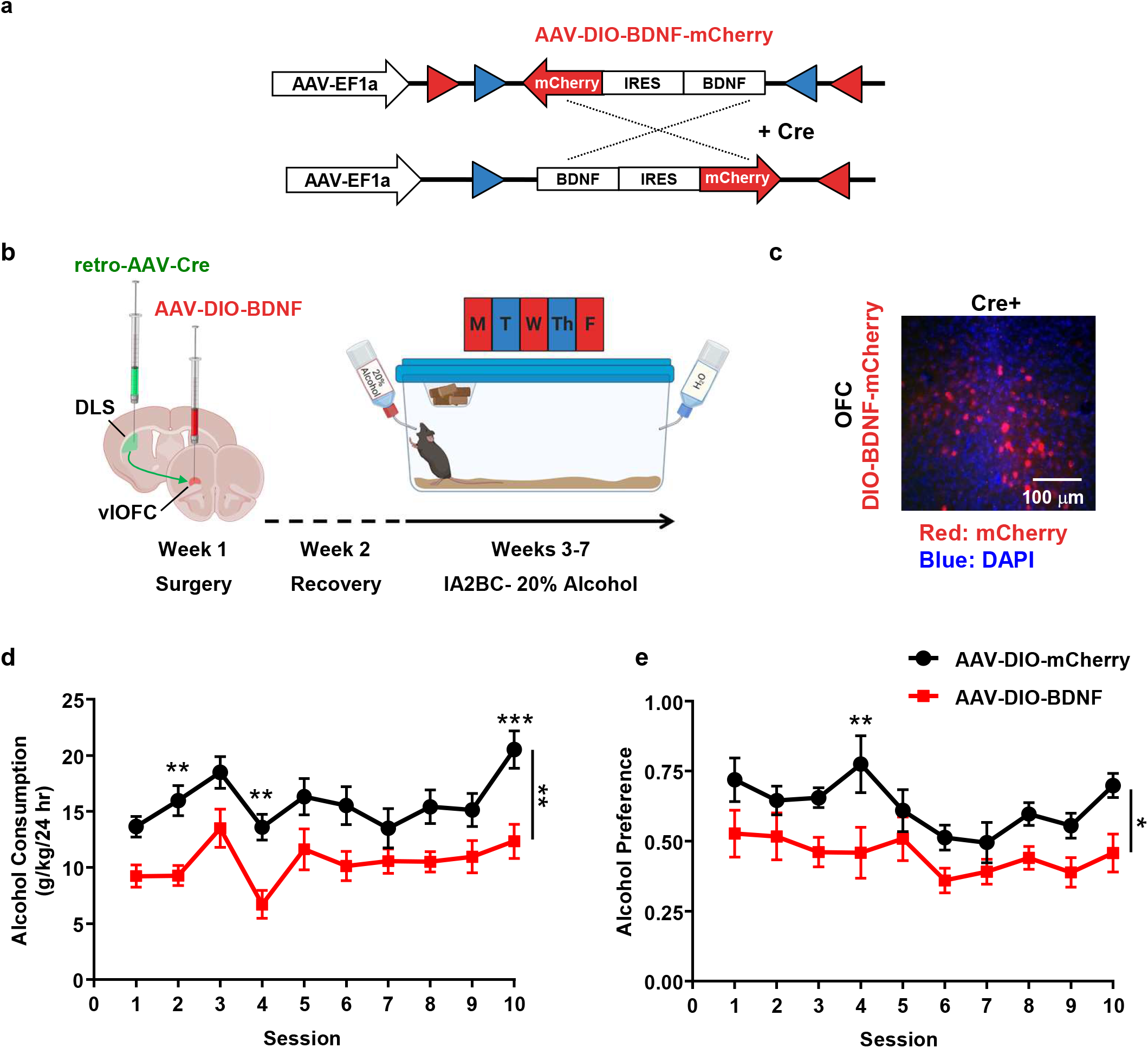
Pathway specific overexpression of BDNF in vlOFC neurons which project to the DLS gates alcohol intake. (**a**) Schematic of the AAV-DIO-BDNF-mCherry construct injected into the vlOFC of test mice. Co-expression of AAV-Cre recombinase and AAV-DIO-BDNF-mCherry leads to an inversion of the BDNF-IRES-mCherry cassette, allowing BDNF and mCherry to be expressed. (**b**) Timeline of experiments: AAV-DIO-BDNF virus or AAV-DIO-mCherry was stereotaxically injected into the vlOFC and a retrograde AAV-Cre viral construct was infused into the DLS. After one week of recovery, mice underwent 10 sessions of IA20%-2BC in the home cage and alcohol intake was recorded at the 24 hour time point. (**c**) High magnification (20X) image depicting mCherry expression in vlOFC cells co-infected with Cre (via retrograde transport from the DLS) and AAV-DIO-BDNF. Red cells indicate co-expression of the Cre and AAV-DIO-BDNF, confirming the overexpression of BDNF in vlOFC neurons projecting to the DLS. (**d**) Overexpressed in vlOFC to DLS circuitry maintains alcohol intake in moderation (Two-Way RM ANOVA, main effect of circuit specific BDNF overexpression, F _(1,13)_ =15.86, ^**^p = 0.001, main effect of session F_(9,117)_ = 6.08, ^***^p < 0.001, and no interaction of virus and session, p > 0.05. Bonferroni post-hoc testing indicates a significant difference between conditions on sessions 2, 4, and 10, ^**^p < 0.01 and ^***^p < 0.001). (**e**) Overexpressed in vlOFC to DLS circuitry reduces alcohol preference (Two-Way RM ANOVA, main effect of circuit specific BDNF overexpression, F _(1,13)_ =8.82, ^*^p = 0.01, main effect of session F_(9,117)_ = 3.40, ^***^p < 0.001, and no interaction, p > 0.05). Bonferroni post-hoc testing indicates a significant difference between conditions on session 4, ^##^p < 0.01), while those receiving control infusions displayed high alcohol preference. AAV-DIO-BDNF n = 8, AAV-DIO-mCherry n = 7.

First, we confirmed that the AAV-DIO-BDNF-mCherry construct was functional by injecting it into the vlOFC of CamKIIα-Cre mice, which express Cre recombinase in cortical glutamatergic neurons, or in wild type C57Bl6/J mice, and examined mCherry fluorescence. We observed a high number of mCherry-positive cells in the vlOFC of CamKIIα-Cre mice that received the AAV-DIO-BDNF-mCherry virus and no mCherry expression in wild type mice that received the same injection. (**Supplemental Figure 4a**). This shows that mCherry and BDNF are only overexpressed in cells that co-express Cre recombinase. Next, to overexpress BDNF only in the vlOFC-DLS pathway, we infused a retrogradely-transported AAV5-Cre-GFP virus into the DLS and AAV-DIO-BDNF-mCherry virus into the vlOFC (**Figure 5b**). We detected a robust retrograde transport of Cre-GFP in the striatum (**Supplemental Figure 4b, bottom panel**), and a high number of mCherry-positive cells in the vlOFC (**Figure 5c, Supplemental Figure 4b**), suggesting that this circuit-specific strategy is an effective approach to overexpressing BDNF in DLS-projecting vlOFC neurons.

One week after infusion, mice were subjected to the IA20%-2BC paradigm, and alcohol and water intake were monitored (**Timeline Figure 5b**). Control animals received AAV-Cre in the DLS and AAV-DIO-mCherry in the vlOFC. As shown in **Figures 5d-e**, overexpression of BDNF in the vlOFC-DLS pathway resulted in a significant decrease in both alcohol consumption and preference over ten sessions. We observed a counterbalanced significant increase in water consumption in mice overexpressing BDNF (**Supplemental Figure 5a**), resulting in no change in total fluid consumption between these experimental groups (**Supplemental Figure 5b**). These results suggest a role for BDNF in vlOFC neurons projecting to the DLS are responsible for keeping alcohol intake in moderation.

## Discussion

Here, we show that BDNF in the mouse vlOFC keeps alcohol drinking in moderation and decreases alcohol seeking behaviors but does not affect saccharin consumption. We further demonstrate that BDNF in DLS-projecting vlOFC neurons gates the level of alcohol consumption. In contrast, we show that BDNF in the motor cortex, which also sends projections to the DLS [25], does not affect home cage alcohol drinking.

The OFC plays roles in decision making and reward learning [26-29], and in forming associations between stimuli and their outcomes [30-32]. Furthermore, Hernandez et al. showed that alcohol seeking activates neurons in the rat OFC, and the level of activation depends on individual preference for alcohol [33]. Therefore, it is possible that BDNF in the vlOFC regulates alcohol consumption and preference by influencing decision making related to alcohol seeking behaviors. Gourley and colleagues reported that BDNF in the OFC is necessary for proper goal-directed response selection for food and/or cocaine rewards following reward devaluation [32,34,35]. Therefore, BDNF in the vlOFC may gate alcohol consumption and seeking behavior by promoting goal-directed alcohol seeking over habitual drinking, which should be addressed in future studies.

The rodent OFC is a complex brain region containing several functionally distinct subregions, with different neuronal populations extending axonal projections throughout the brain [28]. We recently showed that the vlOFC but not medial OFC (mOFC) sends projections to the DLS [25]. In rodents, primates and humans, the vlOFC is an anatomically separate subregion, with distinct corticostriatal circuits linked to specific behavioral functions [28,36]. In rodents and primates for instance, the lateral OFC/vlOFC has been shown to play a larger role in regulating goal-directed behaviors [30,37], behavioral flexibility [38], and relationships between stimuli or actions and their respective outcomes [39]. Future studies to determine whether BDNF also plays a role in regulating alcohol seeking and consumption in other OFC subregions and their projections are warranted.

We found that overexpression of BDNF in vlOFC to DLS circuit moderates the level of alcohol intake and preference. The majority of BDNF activating TrkB signaling in the striatum originates from the cerebral cortex [22-24]. BDNF transduces its signal in part via TrkB-dependent activation of ERK1/2 [40,41], and we recently found that overexpression of BDNF in the vlOFC activates ERK1/2 signaling in the DLS [25]. In addition, we previously showed that intra-DLS infusion of recombinant BDNF reduces rat operant self-administration of alcohol through a mechanism that requires ERK1/2 [13]. Together, these data suggest that BDNF produced in the vlOFC is transported to terminals in the DLS where it activates TrkB signaling, which in turn gates alcohol intake.

The results herein further highlight the importance of the vlOFC to DLS projection in regulating alcohol drinking. Within the DLS, it is also likely that BDNF signaling via TrkB is differentially regulated depending on target cell type, such as dopamine D1 versus D2 receptor-expressing medium spiny neurons [42-44]. Furthermore, the vlOFC and lOFC extend projections to several other brain regions, including the hippocampus, substantia nigra, ventral tegmental area, and several cortical subregions [28,45] and work from Gourley and colleagues, suggests that the vlOFC projections to the ventrolateral striatum, regulate goal-directed food and cocaine seeking [32]. Thus, we cannot exclude the possibility that BDNF originating from the vlOFC activates TrkB signaling in one or more of these brain regions and may also play a role in alcohol intake.

Overall, these results highlight the importance of a little-studied projection in regulating alcohol intake, additional studies will need to examine its potential function in regard to other rewarding substances. Our data provide further insight into mechanisms that can protect against escalating alcohol consumption.

## Funding and Disclosure

This research was supported by the National Institute of Alcohol Abuse and Alcoholism F32AA028422 (JJM) and R37AA01684 (DR). The authors have no conflicts of interest to disclose.

## Author Contributions

JJM designed the study, conducted experiments, analyzed data, and wrote the manuscript. SAS designed the study and conducted experiments. YE and KP conducted experiments. DR designed the study and wrote the manuscript.

Figures were created in part using BioRender.

## Notes

### Competing Interest Statement

The authors have declared no competing interest.

