## Supplemental Figure Legends for "BRAIN-DERIVED NEUROTROPHIC FACTOR IN AN ORBITOFRONTAL CORTICAL-DORSOLATERAL STRIATAL CIRCUIT GATES ALCOHOL CONSUMPTION"

**Supplemental Figure 1. Water and total fluid consumption of mice infected with AAV-BDNF in the vlOFC**

No significant difference in water consumption (**a**) or total fluid (**b**) in in mice infected with AAV-BDNF mice versus mice infected with AAV-GFP. (**a**) Water intake (Two-Way ANOVA, no main effect of BDNF overexpression, $p < 0.05$, session, $p > 0.05$, or interaction, $p > 0.05$) (**b**) Fluid consumption (Two-Way ANOVA, main effect of session, $^{*}p < 0.05$, no main effect of BDNF overexpression, $p < 0.05$, or interaction, $p > 0.05$). AAV-BDNF n = 10, AAV-GFP n = 10.

**Supplemental Figure 2. Home cage water seeking with BDNF overexpression in the vlOFC**

(**a**) No significant difference between mice that received AAV-BDNF in the vlOFC and underwent 5 weeks of IA20%-2BC in time spent licking a water bottle during the first 2 minutes of exposure. Mean time licking is presented with individual data points representing individual animals. (Unpaired t Test, t = 0.06231, df = 15, p (two-tailed) > 0.05). (**b**) Representative traces of one AAV-BDNF and one AAV-GFP mouse's water bottle lick time ratio (the amount of time each mouse spent in contact with the bottle sipper tube per second) during a two-minute alcohol seeking test. (**c**) Latency to first lick of the water bottle. Data is presented as the mean amount of time it took for mice from each group to initiate licking of the water bottle (maximum value = 120 seconds). (Unpaired t Test, t = 2.566, df = 18, p (two-tailed) < 0.05). (**d**) Latency to last lick of the water bottle. Data is presented as the mean amount of time from the final lick of the water bottle to the end of the session for mice from each group (maximum value = 120 seconds). (Unpaired t Test, t = 2.625, df = 18, p (two-tailed) < 0.05). AAV-BDNF n = 10, AAV-GFP n = 10.

**Supplemental Figure 3. BDNF overexpression in the motor cortex does not alter water or total fluid consumption**

(**a**) Animals which received AAV-BDNF or AAV-GFP in the mCtx did not significantly differ in levels of water consumption (Two-Way RM ANOVA, no main effect of BDNF overexpression, $p > 0.05$, interaction, $p > 0.05$, or session $p > 0.05$). (**b**) Total fluid consumption was similar in mice infected with AAV-BDNF or AAV-GFP in the mCtx (Two-Way RM ANOVA, no main effect of BDNF overexpression, $p > 0.05$, interaction, $p > 0.05$, or session $p > 0.05$). AAV-BDNF $n = 7$, AAV-GFP $n = 7$.

**Supplemental Figure 4. Confirmation of AAV-DIO-BDNF-mCherry virus expression**

(**a**) CaMKIIα-Cre mice received injections of AAV-DIO-BDNF-mCherry in the vlOFC to confirm targeting and viral infection (*Top*). Left image is a 5X magnification image of the prefrontal cortex. Right panel is a 20X magnification image of the area of the image in the left panel outlined in white, demonstrating mCherry expression in vlOFC neurons co-expressing Cre and AAV-DIO-BDNF-mCherry. C57Bl6/J mice received AAV-DIO-BDNF-mCherry injections in the vlOFC to confirm the necessity of Cre recombinase co-expression for the construct to be expressed (*Bottom*). 5X image of the prefrontal cortex showing no mCherry expression in neurons only expressing AAV-DIO-BDNF-mCherry (no Cre) (**b**) Confirmation of circuit specificity following injection of retrograde AAV-Cre-GFP in the DLS and AAV-DIO-BDNF-mCherry in the vlOFC. Top left image is a 5X magnification image of the prefrontal cortex. Top right panel is a 20X magnification image of the area of the image in the left panel outlined in white, demonstrating mCherry expression in vlOFC neurons co-expressing Cre recombinase and thus confirming projections to the DLS. Bottom left panel is a 5X image displaying GFP expression in the striatum due to retrograde transport of the AAV-Cre-GFP virus injected into the DLS.

**Supplemental Figure 5. Water and total fluid consumption with circuit-specific BDNF overexpression in the vlOFC-DLS circuit**

(**a**) Animals that overexpressed BDNF in a pathway-specific fashion drank significantly more water during IA20%-2BC (Two-Way RM ANOVA, main effect of circuit specific BDNF overexpression, $F_{(9, 117)}$ =0.6966, $^{*}p < 0.05$, main effect of session $F_{(9,117)} = 6.619$, $^{***}p < 0.001$, and no interaction of virus and session, $p > 0.05$) (**e**) but no significant change in total fluid consumption (Two-Way RM ANOVA, main effect of session $F_{(9,117)} = 0.5397$, $^{***}p < 0.001$, and no main effect of circuit specific BDNF overexpression or interaction, $p > 0.05$). AAV-DIO-BDNF n = 8, AAV-DIO-mCherry n = 7
