## Supplemental Figures for "BRAIN-DERIVED NEUROTROPHIC FACTOR IN AN ORBITOFRONTAL CORTICAL-DORSOLATERAL STRIATAL CIRCUIT GATES ALCOHOL CONSUMPTION"

**Supplemental Figure 1**

### Water and total fluid consumption of mice infected with AAV-BDNF in the vlOFC

a

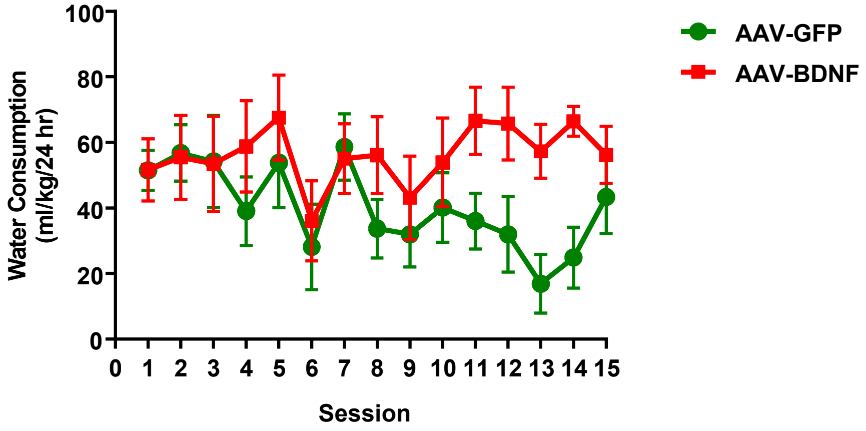


b

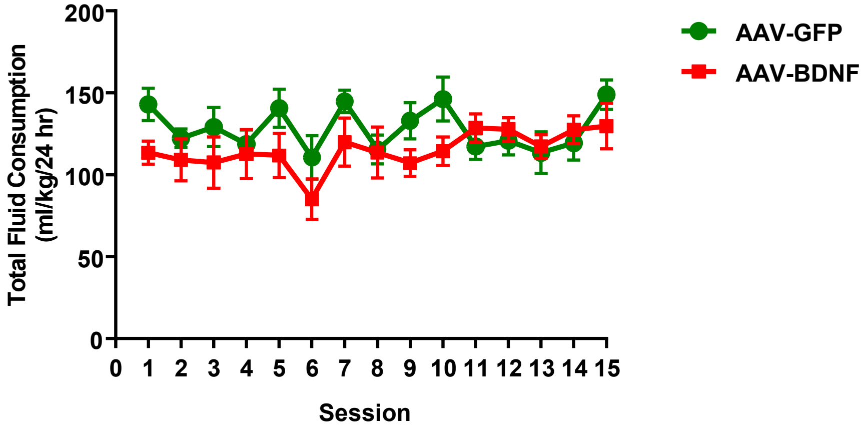

**Supplemental Figure 2**

**Home cage water seeking with BDNF overexpression in the vlOFC**

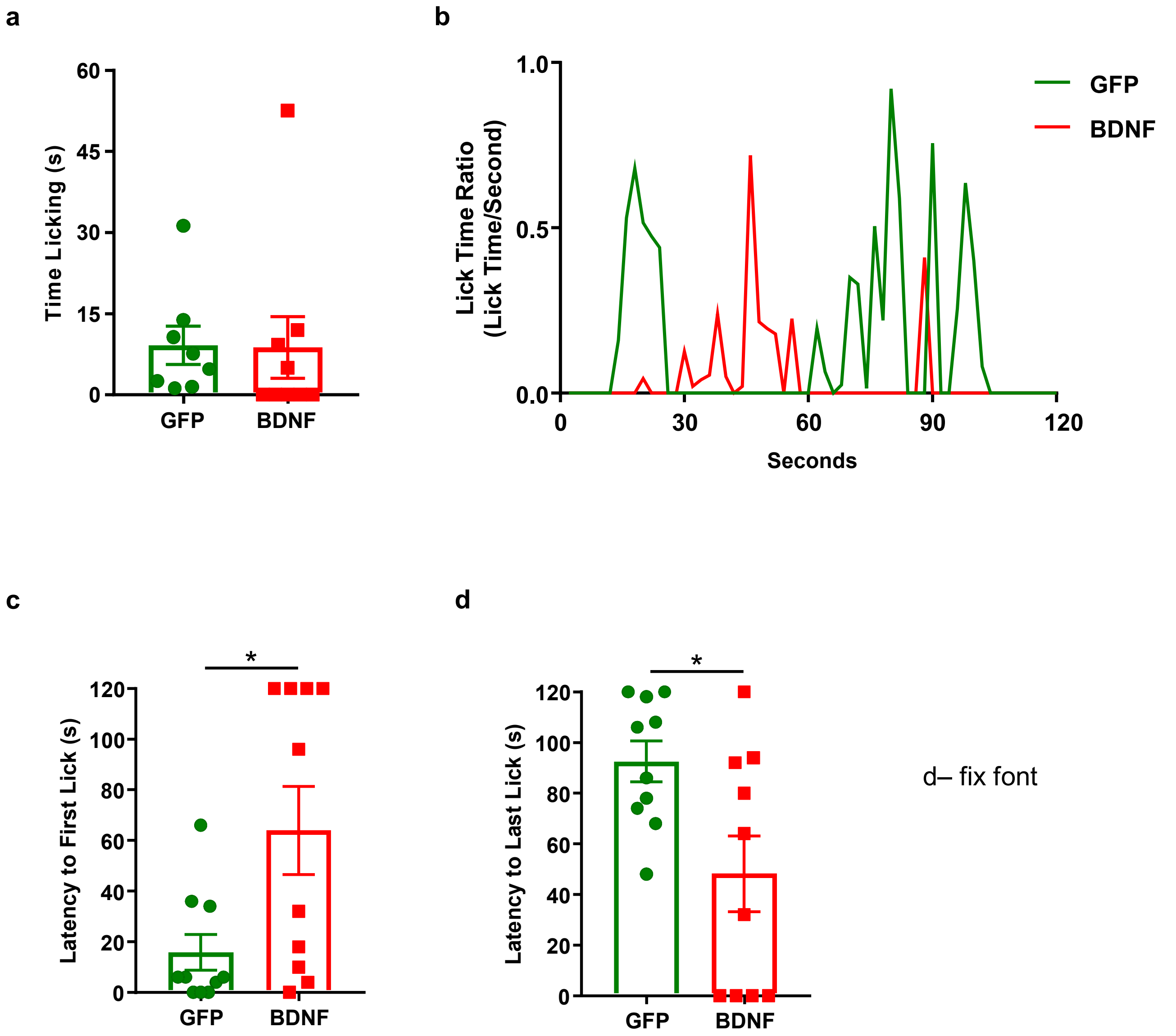


d– fix font

### Supplemental Figure 3

**BDNF overexpression in the motor cortex does not alter water or total fluid consumption**

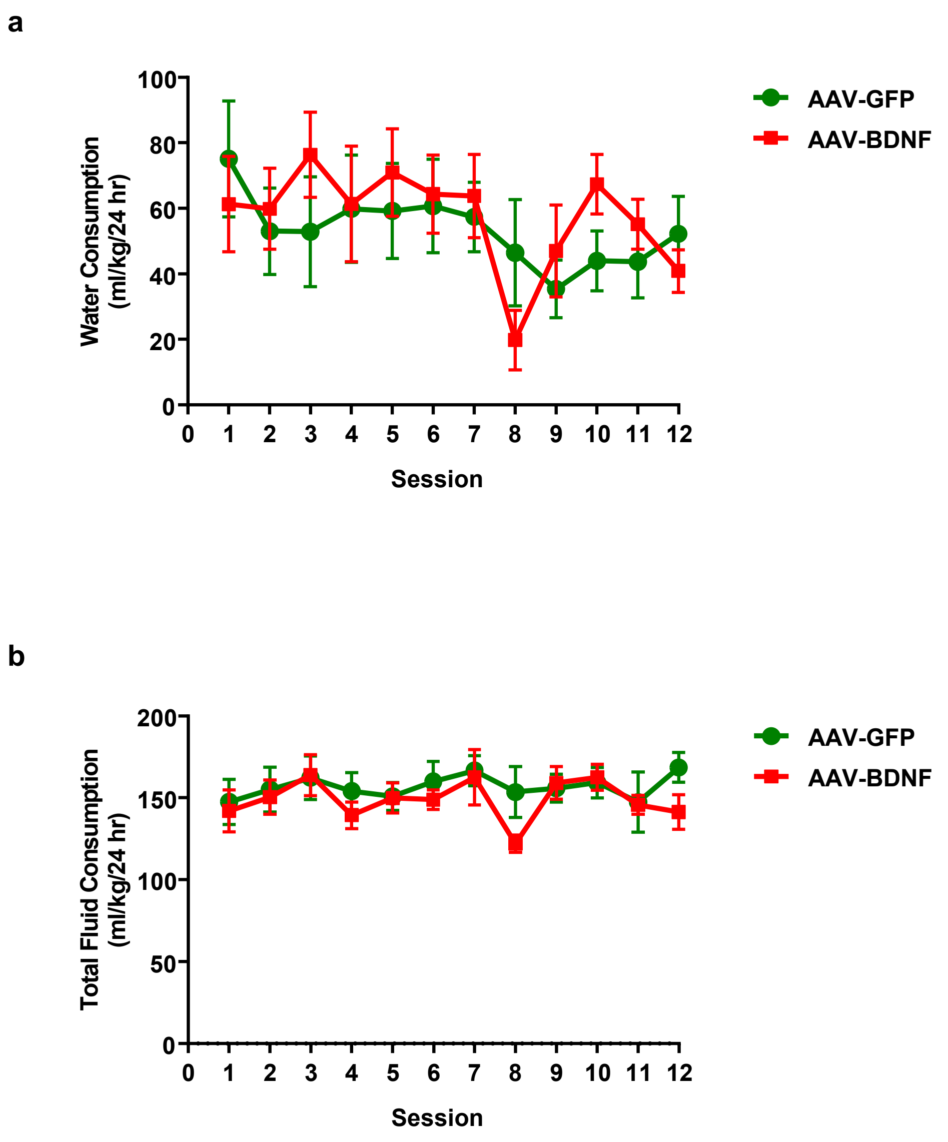

### Supplemental Figure 4

#### Confirmation of AAV-DIO-BDNF-mCherry virus expression

**a**

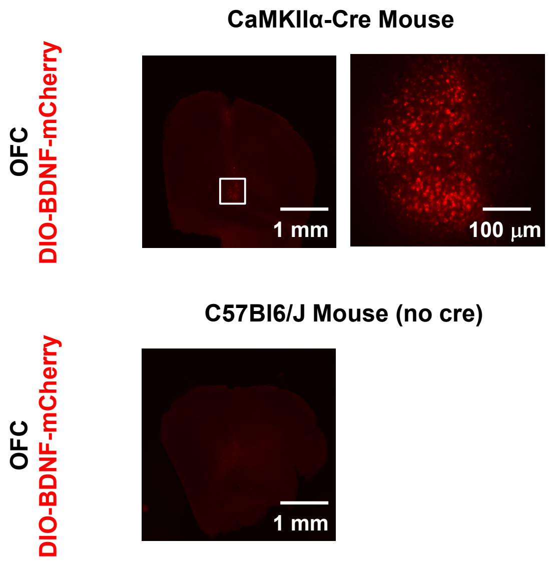


**b**

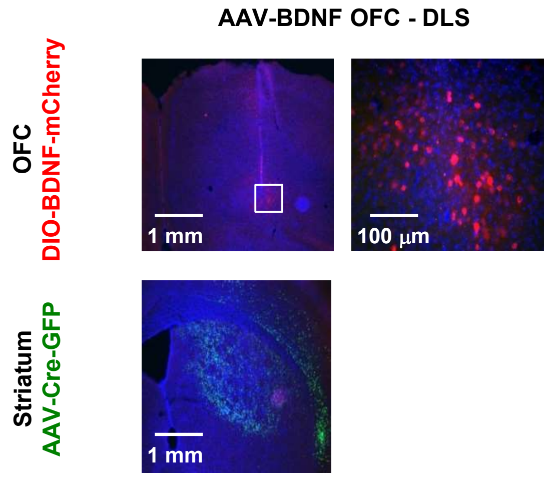

**Supplemental Figure 5**

### Water and total fluid consumption with circuit-specific BDNF overexpression in the vlOFC-DLS circuit

**a**

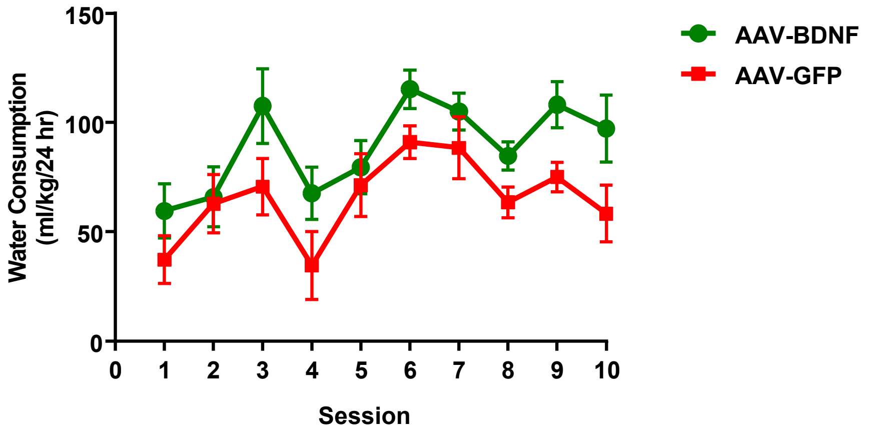


**b**

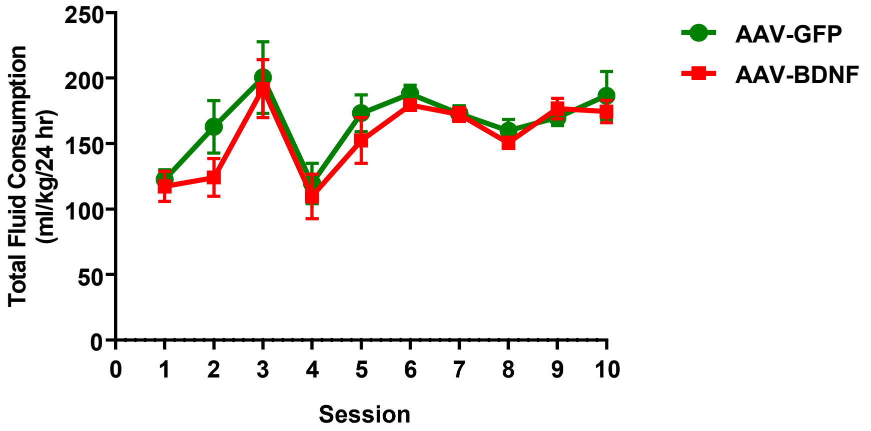
